# From Nutraceutical to Molecular Targets: Antidiabetic Effects of *Spirulina maxima* Validated by In Vivo and In Silico Studies

**DOI:** 10.64898/2025.12.27.696639

**Authors:** Anish Zacharia, Senthil kumar Subramani, Pratibha Chauhan, Mula Kameshwara Rao, Prakash Singh Bisen

## Abstract

**Background:** *Spirulina maxima* is a cyanobacterium, which has been a staple food in Mexico since 16^th^ century. Presently this nutraceutical alga has been explored for its antioxidant, anticancer, hepatoprotective and antilipemic properties. The present study is focused on the antihyperglycemic effect of *S.maxima* in streptozotocin induced diabetes mellitus. The molecular characterization of *S. maxima* and the anti-diabetic targets are deduced from *in-silico* analysis.

**Methods:** The anti-hyperglycemic potential of *S. maxima* was evaluated in male Wistar rats injected with streptozotocin (45 mg/kg (i.p)). The Animals were grouped based on the blood levels into Normal control (n=6); Normal control administered with *S. maxima* (n=6), Diabetic control, Diabetic group treated with *S. maxima*(n=6),, the diabetic group treated with Glibenclamide (600µg/kg b.w.) (n=6), *S. maxima* (500mg/kg) was administrated orally for 15 days, and the variations in blood glucose levels along with the lipid levels and the antioxidant status were determined.The lipids content of *S. maxima* was characterized through GCMS. The lipids identified were docked *in-silico* with AMPK molecule and PPARγ, potential targets for antidiabetic drugs.

**Results and conclusions:** *S. maxima* administration in diabetic rats significantly reduced the blood glucose levels by 62.76% while the cholesterol, triglycerides, and VLDL level were marginally reduced. The HDL levels were significantly enhanced by 32.4%. The antioxidant markers such as catalase, SOD and GSH levels, were elevated by 13.8, 25 and 7.47%. The TBARS levels were reduced up to 15.1%. The functional group analysis showed characteristic vibrations of polysaccharides proteins, lipids and aromatic compounds. The GCMS characterization of *S. maxima* showed the presence of C_8_, C_9_ and C_13_ lipids at a concentration less than 1% of total lipids, while C_5_, C_11_, C_16_ and C_20_ lipids were 1-10% of total lipid content while C_6_, C_10_, C_15_ and C_22_ lipids were above 10% of total lipids. The docking results revealed the binding affinity of C_5_, C_6_, C_10_, C_11_ and C_18_ with regulatory gamma subunit of AMPK while C_5_, C_6_ and C_11_ lipids had the binding affinity with PPAR-γ receptor. The study revealed, the antidiabetic potential of *S. maxima*, and hypothesized AMPK and PPAR-γ as a potential target in maintaining blood glucose homeostasis.

## 1. Introduction

This work is dedicated to the memory of Late Prof. Prasad, a distinguished researcher in the field of clinical diabetes at Jiwaji University. His pioneering contributions to diabetes research, combined with his unwavering commitment to academic excellence and mentorship, have left a lasting impact on both the scientific community and his students. Prof. Prasad’s insightful guidance and passion for translational research continue to inspire ongoing efforts to understand and manage metabolic disorders.

Diabetes mellitus is a chronic metabolic disorder associated with the abnormalities in carbohydrate, lipid metabolisms American Diabetes Association (2009). This metabolic disorder is responsible for the high rates of mortality in both developed and developing countries. Diabetes in association with oxidative stress and inflammation can lead to microvascular and macrovascular complications. Hence there is always a need of drug which addresses the diabetes mellitus and its associated disorder. *Spirulina* is blue green algae known for its nutritional value and therapeutic potential (Khan et al., 2005; Kulshreshtha et al., 2008). Presently various therapeutically active compounds have been identified in this organism.

Spirulina administration has been proved effective in preventing the hyperlipidemia (Torres-Duran et al., 1998, 2007), fatty liver condition (Blé-Castillo et al., 2002) and hypertension (Ferreira-Hermosillo et al., 20l0) in animals and humans. *S. maxima* was reported to be a rich source of antioxidant molecules like tocopherols, β-carotene (Miranda et al., 1998). The *S. maxima was* also reported to normalize the fructose induced hyperglycemia in rats (Jarouliya et al., 2012). Spirulina consumption showed no adverse or toxic effects and has been confirmed in several studies. (Yoshino et al., 1980; Krishnakumari et al., 1982; Chamorro et al., 1996; Salazar et al., 1996). The present study was intended to evaluate the antidiabetic potential of S*. maxima* in Streptozotacin induced diabetes mellitus in a rat model. Attempts were made to characterize the lipid metabolites of *S. maxima* through GCMS. Furthermore, function of the characterized molecule was envisaged through molecular docking approaches.

## 2.0 Material & Methods

### Animals

Randomly bred healthy male rats (Rattus norvegicus; Wistar strain; age: 10-14 wk; weight: 180-220g) were obtained from the animal facility of Defence Research & Development Establishment, Gwalior (MP). The rats were housed in polypropylene cages, and were maintained on standard pellet diet (M/s Ashirwad Industry, Mohali, India) and water *ad libitum*. After randomization into various groups and before initiation of the experiment, the rats were acclimatized for a period of seven days to laboratory conditions [temperature (25 ± 2°C) and 12 h light/dark cycle]. All the chemicals used in the experiments were of analytical grade, purchased from E. Merck India Limited (Mumbai, India).

### 2.1 Culturing of Spirulina maxima

The pure culture of *S. maxima* was obtained from National Facility for Blue Green Algae, Indian Agriculture Research Institute, New Delhi. The culture was grown in Zarrouk’s medium (pH 9.0) under controlled conditions of light and temperature (5000 lux, 12/12 h light/dark, at 30°C) in an airlift Photobioreactor. The cultures were maintained for 12 days and harvested at exponential phase of growth. Biomass was filtered through a screen filter of pore size 305 nm. The Spirulina mat was collected in the drying trays and dried at 40°C for 12 h.

#### Dose Optimization studies by OGTT method

The dose optimization of *S. maxima* was by oral glucose tolerance tests. The animals were grouped based on their fasting blood glucose levels. The study was performed in normal animals (n=10). The rats were fasted O/N, and their fasting blood glucose levels were monitored by ACCUCHECK glucometer and followed by oral administration of glucose 2g/kg b.w. The different doses of *S. maxima* (100mg/kg, 250 mg/kg and 500mg/kg) along with allopathy drug Glibenclamide (600μg/kg) were administered to normal rats and changes in blood glucose levels at every 30 min interval for two hours were observed

### 2.2. Induction of hyperglycaemia

Hyperglycemia was induced in male Wistar rats by intraperitoneal administration of Streptozotocin (45 mg/kg i.p), and the animals were divided into five groups of five each and treated as below Group I: Healthy rats (Normal control): Normal rats without any treatments; Group II: Normal rats administered orally with *S. maxima* (500 mg/kg); Group III: Diabetic control; Group IV: Hyperglycaemic rats administered *S. maxima* (500 mg/kg); Group V: Hyperglycaemic rats administered orally with Glibenclamides (600µg/kg b.w).

#### Blood collection

The animals were kept on overnight fasting, and then the blood samples (0.5-1.0 ml) were collected from retro orbital plexus under mild ether anaesthesia once before and at 10 day intervals during the course of study. Plasma was separated, aliquoted and used for analyses of blood glucose, triglyceride and cholesterol, remaining plasma sample was assayed for HDL cholesterol, LDL, VLDL All estimations were carried out on freshly collected blood/plasma samples through the diagnostic kits. The blood glucose levels were estimated by GOD-POD method (Trinder, 1969). The triglycerides were estimated byGPO-POD method (Wood et al., 1998). The Cholesterol levels were estimated through CHOD-PAP method (Deeg et al., 1983). The HDL cholesterol levels were estimated by Phosphotungstate method (Lopes-virella, 1977). The LDL and VLDL levels were calculated by Friedewald’s equation.

### 2.3 Sample preparation for oxidative stress markers

The oxidative stress markers such as GSH was assayed in whole blood and catalase, superoxide dismutase and TBARS activity were assayed in hemolysate.

The packed cell volume (PCV) or hemolysate was prepared after the removal of the plasma and the buffy coat from whole blood cells by centrifugation at 2000 rpmx10minx4°C. The red cells were further washed thrice with chilled normal saline (0.9%) to get the PCV. From this PCV, hemolysate(s) for further estimations were prepared as follows. For the determination of CAT and TBARS, the hemolysate was prepared by making 5% PCV solution in chilled distilled water.

#### 2.3.1. TBARS Levels

The TBARS levels were determined according to the method of Ohkawa et al., (1979). The Malonaldehyde (MDA), a decomposition product of lipid hydro peroxide was used as an indicator of oxidative damage to cells and tissues. In this method, the test sample was heated with thiobarbituric acid (TBA) under acidic conditions resulting in the development of pink colour. A small amount of MDA is produced during peroxidation which reacts with the TBA in this test generated a colored product, and the absorbance was read at 532nm using spectrophotometer. The amount of TBARS was calculated using a molar extinction coefficient of 1.56 x10^-5^ mole/mm and the values are expressed as nmole/mg protein.

#### 2.3.2. Catalase activity

The Catalase activity was assessed according to the method of Sinha (1972).

This method is based on the fact that the dichromate in acetic acid is reduced to chromic acetate when heated in the presence of H_2_O_2_, with the formation of perchromic acid as an unstable intermediate. The chromic acetate thus produced was measured colorimetrically at 560-630 nm. The catalase activity was expressed in μmole/min/mg protein.

##### Sample preparation for SOD activity

For the determination of SOD, the remaining red cells were hemolyzed by adding distilled water. The lipids were removed by using chloroform-ethanol extraction. For this, the hemolysate was diluted four times with ice-cold distilled water. To 4 ml hemolysate, 1ml of ethanol and 0.6ml of chloroform were added sequentially with continuous shaking for 1 min. The preparation was subjected to centrifugation at 3000rpmx10minx4°C. The supernatant was used for the estimation of SOD.

#### 2.3.3. SOD activity

The SOD activity was assessed according to the method of Winterbourn et al.,(1975). This method is based on the formation of NADH-phenazine methosulphate-nitro blue tetrazolium formazan complex. Acetic acid was used to arrest the formazan formation. The colour intensity of the chromogen was measured at 560 nm. The SOD activity was expressed in units/min/mg protein.

#### 2.3.4. GSH activity

The GSH activity was assessed according to the method of Ellmann et al., (1979). This method is based on the fact that DTNB {5,5-dithio bis (2-nitro benzoic acid)} is reduced by sulfhydryl group (SH), present in reduced glutathione to form one mole of 2-nitro −5-mercaptobenzoic acid per mole of SH. The reaction is as follows. The nitromercaptobenzoic acid anion has an intense yellow colour and is used to measure SH groups. Development of maximum colour intensity occurred immediately after the addition of DTNB, however, absorbance decreases rapidly with time. The absorbance was measured at 412 nm. The GSH levels were expressed in mg/ml.

#### 2.3.5. Total Protein content

Total Protein content was assessed according to the method of Lowry et al., (1951). Protein reacts with the Folin-Ciocalteau reagent to give a colored complex. The colour so formed is due to the reaction of the alkaline copper with the protein as in the biuret test and the reduction of phosphomolybdate by tyrosine and tryptophan present in the protein. The intensity of color depends on the amount of these aromatic amino acids present and will thus vary for different proteins. Optical density was measured spectrophotometrically at 660 nm against the blank. The amount of total cellular protein was expressed as mg/ml

### 2.4. GCMS profiling of lipid content of *S. maxima*

The metabolite profile of *S. maxima* was analyzed by GCMS. The dried S. maximum (1gm dry weight) was extracted in 50 ml of chloroform and methanol at proportion of 2:1. The material was kept on a magnetic stirrer for 4 hours at room temperature. The extract was filtered using whattmann no1 filter paper. Repeat the experiment and the extract was collected and half the volume of 0.9% Nacl was added. The organic phase was carefully separated and evaporated to dryness.

#### 2.4.1. Derivatization of the *S. maxima* lipids

BF3-Methanol, 10% w/w (10% boron trifluoride in methanol) is particularly useful for preparing methyl esters of carboxylic acids and esters (C8-C24 chain length). The sample and the reagent (2ml BF3-methanol at a concentration of 10% w/w) were heated in a sealed screw capped vial at 60^0^C for 10 minutes. The analytes were combined with the anhydrous alcohol (methanol) in the presence of an acid catalyst (BF3). In the reaction, the analyte and alcohol molecules are joined with a loss of water. The sample was further cooled and 1ml of water and hexane were added. The reaction vessel was shaken properly for the solubility of esters into the nonpolar solvent. The organic layer was carefully removed and evaporated to dryness. The derivatized lipid sample was reconstituted using diethyl ether and 1µl was injected into GCMS. The concentration of individual lipids in the sample was determined from the calibration of FAME standard lipids ranging from C4-C24. (Supleco).

### 2.5. Receptor Preparation for Docking

The PDB structure of AMPK and PPAR-γ was retrieved from the Research Collaborator for Structural Bioinformatics (RCSB) Protein Data Bank (PDB) (RCSB, www.rcsb.org/). The protein preparation was the first step for molecular docking. The removal of water molecules, metal ions, cofactors, and the addition of charges and hydrogen atoms was done in this step using SPDBV (http://spdbv.vital-it.ch/). Binding Site Prediction Binding sites were characterized by CASTp, Q-Site finder and compared by extensive literature search. By comparing prediction of CASTp algorithm, PASS and Q-Site Finder, best active sites were identified. CASTp method was used to identify and measure the binding sites, active sites, surface structural pockets (accessible), interior cavities (inaccessible), shape (alpha complex and triangulation), area and volume (solvent and molecular accessible surface) of each pocket and cavities of proteins.

#### 2.5.1. Ligand selection and preparation

The structurally homologous lipid molecules of *S. maxima* were obtained from the PubChem compound database. The molecular information was further converted to PDB file through Open Babel software (www.openbabel.org/). Ligand preparation includes addition of hydrogen atoms, neutralization of the charge groups and removal of any miscellaneous structures from the ligand. Prepared and optimized structures of ligand and protein were ultimately used for molecular docking.

#### 2.5.2. Docking

The docking of unliganded PPAR-γ was performed against the different derivatives of lipids retrieved from PubChem compound and docking was performed by using Autodock4 (http://autodock.scripps.edu/wiki/AutoDock4). The Python scripts in MGL tools package were used to analyze the docking results. Python Molecular Viewer (PMV) was used to observe a molecule surface at advanced level or 3D structure.

## Results

### 3.1. Dose optimization studies in normoglycemic rats by OGTT

Oral glucose tolerance test (OGTT) was performed to determine the blood glucose lowering effect of *S. maxima* doses. Different doses of *S. maxima* (100mg/kg, 250 mg/kg, and 500 mg/kg) suppressed the rise in blood glucose level by 28.85%, 37.54% and 46.17%, respectively. Among the three doses of *S. maxima*, 500mg/kg showed the maximum blood glucose reduction (Table 1).

**Table 1:** Effect of *S. maxima* on blood glucose levels in normal wistar rats following oral administration of glucose. **Abbreviation**s: SM – *Spirulina maxima* Units: mg/dl The values were expressed as mean ± SEM of n=10. ^a^p<0.05 (*), ^a^p<0.001 (**) significant compared to normal control group. ^b^p<0.05 (*), ^b^p<0.001(**)significant compared to normal control along with *S.maxima*

| Groups | Blood Glucose levels(mg/dl) |  |  |  |  |
| --- | --- | --- | --- | --- | --- |
|  | 0 min | 30 min | 60 min | 90 min | 120 min |
| Normal control | 78.20 ± 1.56 | 86.80 ± 5.96 | 67.6 ± 1.28 | 91.28 ± 1.47 | 63.6 ± 1.60 |
| Normal (2gm/kg glucose) | 77 ± 1.30 <sup>a**</sup> | 171.12 ± 2.12 <sup>a**</sup> | 161.25 ± 1.83 <sup>aa**</sup> | 153.12 ± 1.97 <sup>a**</sup> | 135.87 ± 0.6 <sup>a**</sup> |
| *S. maxima* (100 mg/kg) | 72 ± 3.17 <sup>a**</sup> | 138 ± 5.45 <sup>a**</sup> | 145.6 ± 6.60 <sup>a**</sup> | 113.66 ± 8.86 <sup>a**</sup> | 96.66 ± 11.35 <sup>a**</sup> |
| *S. maxima* (250 mg/kg) | 81.42 ± 1.61 <sup>a**</sup> | 134.71 ± 8.01 <sup>a**</sup> | 126.28 ± 8.20 <sup>a**</sup> | 95.85 ± 3.24 <sup>a**</sup> | 88.28 ± 2.04 <sup>a**</sup> |
| *S. maxima* (500 mg/kg) | 80.71 ± 1.80 <sup>a**</sup> | 107.57 ± 2.16 <sup>a**</sup> | 91.28 ± 1.47 <sup>a**</sup> | 82.42 ± 0.79 <sup>a**</sup> | 76.71 ± 1.19 <sup>a**</sup> |

#### 3.1.1. Blood glucose lowering effect in diabetic rats

The Fasting blood glucose levels, before diabetes induction seemed to have no obvious difference among any of the groups (Gp I-V). The streptozotocin induction (GP III,IV and V) had elevated Fasting blood glucose levels as compared to their normal controls (P<0.001). The *S. maxima* therapy in normal animals showed no significant change in the FBG levels through out the study period. Subsequently *S. maxima* administration in diabetic animals showed a significant reduction in FBG levels by 62.7% (P<0.005) as compared to the diabetic control group. While no significant difference was observed between *S. maxima* administered diabetic group and Glibenclamide treated groups (Figure 1A).

**Figure 1.**
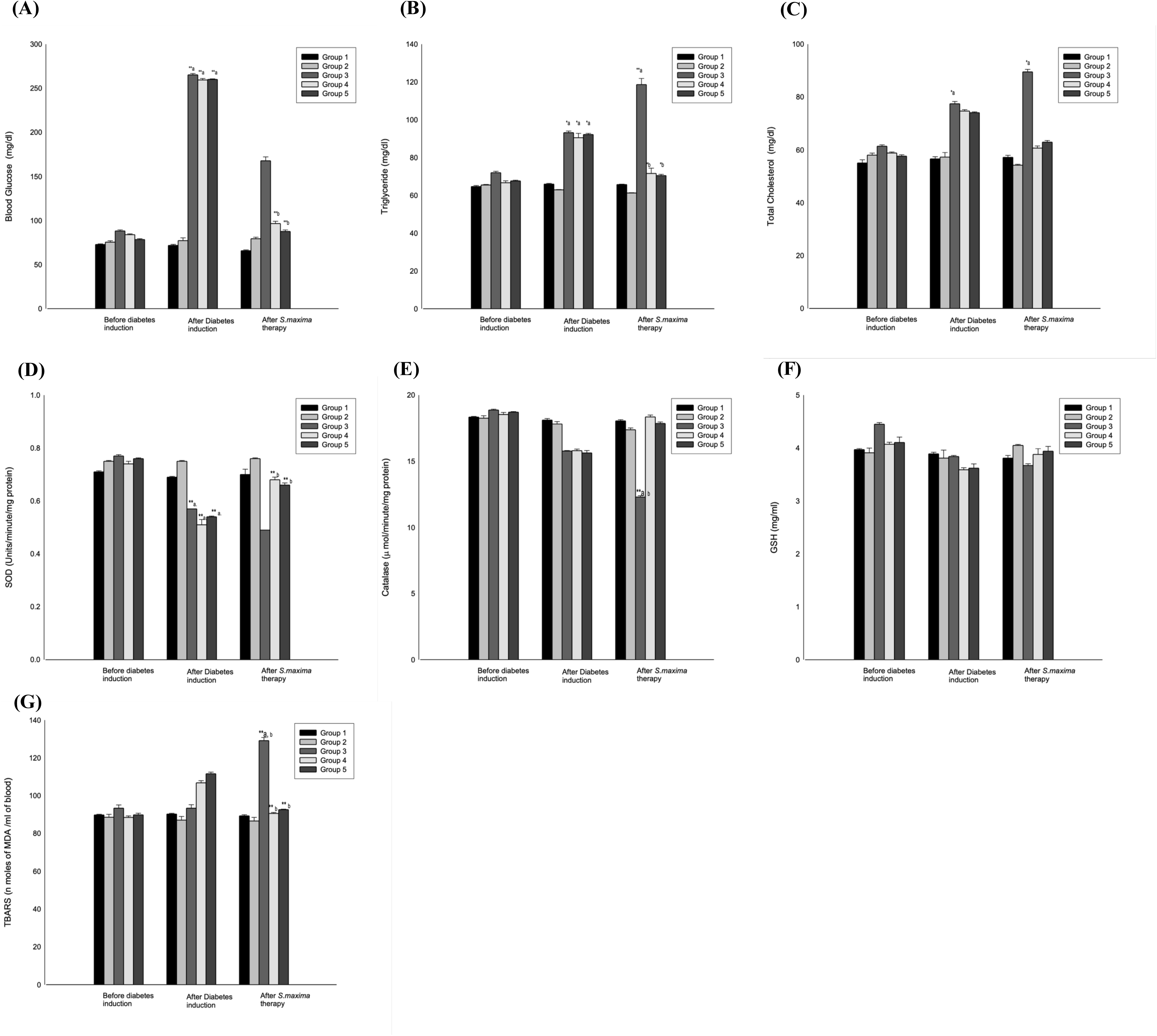
Effect of *Spirulina maxima* on Glycemic Control, Lipid Profile, and Oxidative Stress Markers in Streptozotocin-Induced Diabetic Rats. Effect of *S. maxima* on the (A)Fasting blood glucosse levels (B)Triglycerides (C) Cholesterol (D) SOD (E) Catalase (F) GSH (G) TBARS levels of normal and diabetic rats for 15 days, the values were expressed as Mean + S.E., n=5, Significant change P<0.05(*)b and P<0.001(**)b a as compared after diabetic induction. Significant change P<0.05 and P<0.001(**) a as compared with the before induction Abbreviation: Group 1; Normal control, Group 2; Normal control administered with *S. maxima,* Group 3; Diabetic control, Group 4; diabetic group administered with *S. maxima,* Group 5; Diabetic group administered with glibenclamide. Two-way analysis of variance was determined using statistical software Graphpad Prism Version 5.02.

#### 3.1.2 Lipid regulatory effects in diabetic rats

The STZ induction marginally elevated the cholesterol, triglycerides, VLDL and LDL levels. As summarized in the table, the cholesterol and triglyceride levels were reduced significantly by 18.7% and 20.9% (P<0.05) as compared to diabetic control and normal levels *S. maxima* therapy also reduced the LDL and VLDL levels significantly by 14.38 and 20.9% (Figure 1 B, C) (Table 2-4).

**Table 2:** Effect of *Spirulina maxima* administration on VLDL level in streptozotocin induced hyperglycemia in rats. **Abbreviations:** GP- Group, NC – Normal Control, DC – Diabetic control, TG – Treated Group, GB – Glibenclamide, SM – *Spirulina maxima* **Units:** mg/dl Values are expressed as mean ± SE Significant change P<0.05(*) Significant change P<0.05 and P<0.001(**) Two-way analysis of variance was determined using statistical software Graphpad Prism Version 5.02.

| Groups | Before Diabetic Induction | After Diabetic Induction | After Spirulina Therapy |
| --- | --- | --- | --- |
| GP-I (NC) | 12.93 ± 0.47 | 12.39 ± 0.28 | 13.61±0.303 |
| GP-II (NC)<br>500mg/kg/bw SM | 11.1 ± 0.42 | 12.38 ± 0.39 | 13.88 ± 0.32 |
| GP-III (DC) | 12.1 ± 0.102 | 12.68 ± 0.103 | 10.8 ± 0.136 |
| GP-IV (TG)<br>500mg/kg/bw SM | 12.53 ± 0.38 | 12.3 ± 0.49 | 14.6 ± 0.114 |
| GP-V (TG)<br>600µg/kg/bw GB | 13.33 ± 0.24 | 13.64 ± 0.37 | 17.62 ± 0.86 |

The *S. maxima* administration in diabetic rats for 15 days reduced the cholesterol, triglycerides, VLDL and LDL levels by 18.7, 20.9, 20.9 and 14.38% as compared to the diabetic control (Table 2-4). The *S. maxima* administration in normal animals showed no difference in the total cholesterol, triglycerides, VLDL, and LDL levels. The Glibenclamide treated rats also showed the reduction in cholesterol, triglycerides, VLDL and LDL levels. We have identified that *S. maxima* administration effectively addressed the lipid levels in the diabetic animals. The HDL cholesterol was significantly increased by 24.2% after *S. maxima* therapy as compared to the normal control group (Table 3). Only 3.1% increase was observed in *S. maxima* administered normal animals. The Glibenclamide treated group showed 20.49% increase in HDL content. The present investigation revealed the lipid lowering effect of *S. maxima* in diabetic conditions.

**Table 3:** Effect of *Spirulina maxima* administration on HDL-C level in streptozotocin induced hyperglycemia in rats. **Abbreviations:** GP- Group, NC – Normal Control, DC – Diabetic control, TG – Treated Group, GB – Glibenclamide, SM – *Spirulina maxima* **Units:** mg/dl Values are expressed as mean ± SE Significant change P<0.05(*) Significant change P<0.05 and P<0.001(**) Two-way analysis of variance was determined using statistical software Graphpad Prism Version 5.02.

| Groups | Before Diabetic Induction | After Diabetic Induction | After Spirulina Therapy |
| --- | --- | --- | --- |
| GP-I (NC) | 34.57 ± 0.19 | 36.24 ± 0.21 | 34.09 ± 0.59 |
| GP-II (NC)<br>500mg/kg/bw SM | 35.76 ± 0.37 | 37.87 ± 0.36 | 39.09 ± 0.63 |
| GP-III (DC) | 41.76 ± 1.14 | 31.44 ± 1.62 | 27.06 ± 0.97 |
| GP-IV (TG)<br>500mg/kg/bw SM | 36.66 ± 0.42 | 29.39 ± 0.105 | 38.82 ± 0.52 |
| GP-V (TG)<br>600µg/kg/bw GB | 38.36 ± 0.502 | 30.02 ± 0.25 | 37.76 ± 0.35 |

**Table 4:** Effect of *Spirulina maxima* administration on LDL level in streptozotocin induced hyperglycemia in rats. **Abbreviations:** GP- Group, NC – Normal Control, DC – Diabetic control, TG – Treated Group, GB – Glibenclamide, SM – *Spirulina maxima* **Units:** mg/dl Values are expressed as mean ± SE Significant change P<0.05(*) Significant change P<0.05 and P<0.001(**) Two-way analysis of variance was determined using statistical software Graphpad Prism Version 5.02.

| Groups | Before Diabetic Induction | After Diabetic Induction | After Spirulina Therapy |
| --- | --- | --- | --- |
| GP-I (NC) | 10.48 ± 0.246 | 3.384 ± 0.239 | 4.216 ± 0.119 |
| GP-II (NC)<br>500mg/kg/bw SM | 14.41 ± 0.119 | 5.75 ± 0.832 | 14.21 ± 1.19 |
| GP-III (DC) | 3.038 ± 1.149 | 2.802 ± 1.139 | 1.222 ± 1.170 |
| GP-IV (TG)<br>500mg/kg/bw SM | 9.612 ± 1.188 | 5.024 ± 1.147 | 7.337 ± 1.242 |
| GP-V (TG)<br>600µg/kg/bw GB | 6.711 ± 0.369 | 1.678 ± 0.496 | 2.658 ± 0.499 |

#### 3.1.3 Effect of *S. maxima* on Antioxidant status

Streptozotocin (STZ) involves the generation of nitrogen monoxide (NO) and further resulting in the superoxide O_2_^-^. The systemic levels of antioxidant enzymes that counteract the oxidative stress was reduced. As shown in the Table, the GSH levels in STZ induced group were reduced by 11.7-13.7%, furthermore, the reduction up to 17.5% was observed only in diabetic control group at the end of the investigation. The *S. maxima* administration in diabetic animals enhanced GSH levels by 8.07%. The normal rats administered with *S. maxima* also showed an elevation in GSH levels by 6.29%. The Glibenclamide treated rats enhanced the GSH levels by 8.83% (Figure 2F). Superoxide Dismutase activities were significantly reduced after STZ induction, markedly a reduction range of 25.9-31% was observed after diabetes induction furthermore the reduction up to 36.3% was observed only in diabetic control group at the end of the investigation. The *S. maxima* administration in diabetic animals, enhanced SOD levels significantly by 33.3% (P<0.001) as compared to the diabetic control. While in normal animals, no obvious increase in the levels of SOD was observed after S*. maxima* therapy. The Glibenclamide administration enhanced the SOD levels by 22.2% (Table 2D).

**Figure 2:**
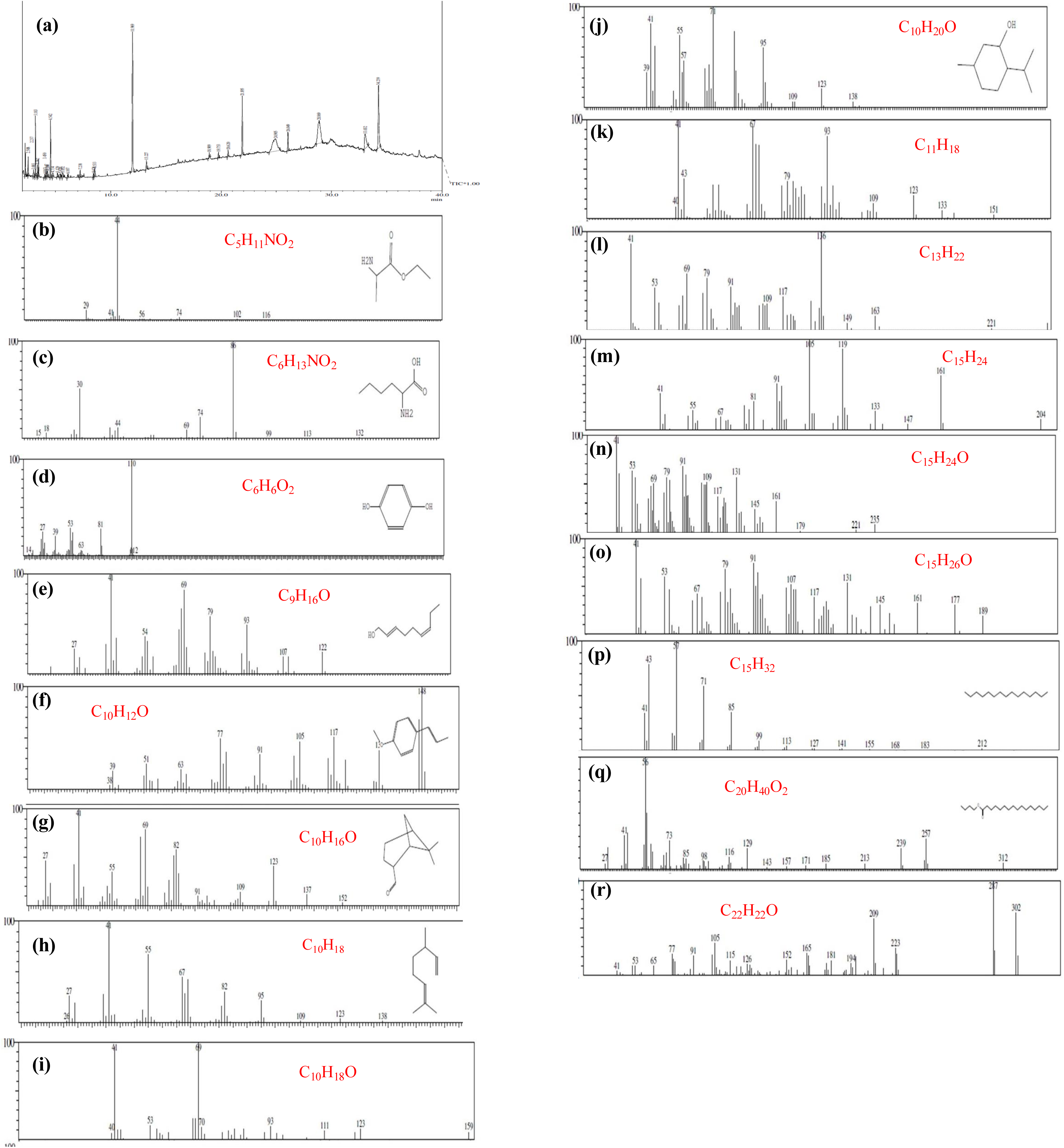
The GCMS spectra of *S. maxima*. The lipid sample of unexposed *S. maxima* were analyzed through GCMS (Shimadzu QP-2010 Plus with Thermal Desorption System TD 20). The sample was eluted through AB inno-wax column (60 m × 0.25 mm id, film thickness 0.25 μm). Helium gas was used as a carrier gas at flow rate of 1.21ml/min. The column oven temperature was maintained at 140.0°C and the injection temperature was maintained at 270^0^C. The ionization temperature was set at 250^0^C-280^0^C. The mass spectra of the analyzed lipids were collected and matched with the published lipid database (Nature lipidomics, wiley 9.0 database, mascot, nist databases. The percentage of each class of lipids present in S. maxima were determined from the standard plot of FAME.

**Figure 2.**
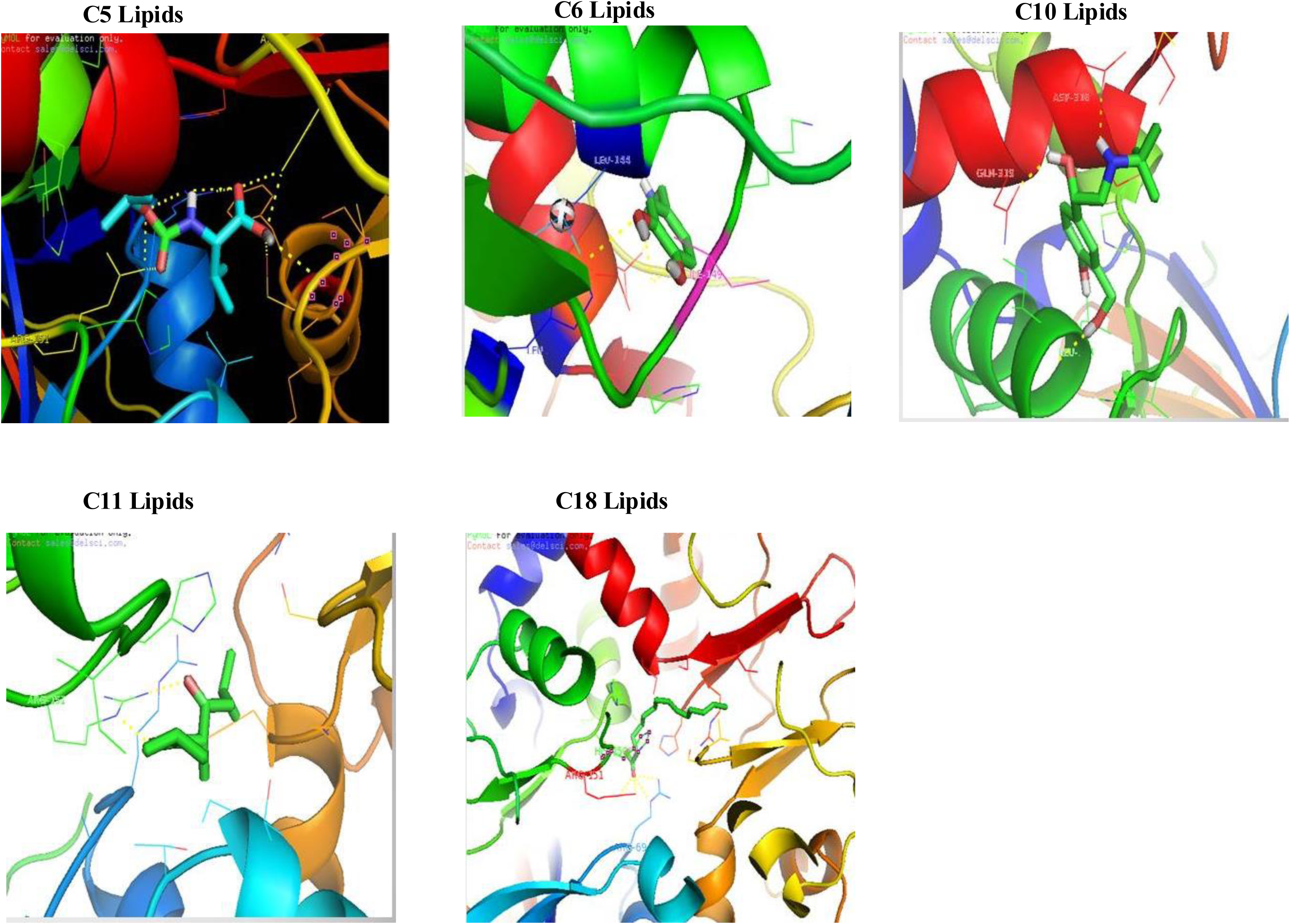
In silico Docking of Bioactive Lipids from *Spirulina maxima* with AMP-Activated Protein Kinase (AMPK)

Catalase activities were significantly reduced after STZ induction, markedly a reduction range of 14.7-16.4% in diabetic groups. Furthermore, the reduction up to 34.8% was observed in diabetic control group at the end of the investigation. The *S. maxima* administration in diabetic animals enhanced the CAT levels significantly by16.13%. While in normal animals, no obvious decrease in the levels of CAT was observed after S*. maxima* therapy. The Glibenclamide administration enhanced the CAT levels by 14.33% (Figure 1E).

TBARS levels were significantly elevated after STZ induction, markedly 20.67-24.27% increase in the TBARS levels were observed in *S. maxima* and Glibenclamide treated group. The Diabetic control group showed elevation in TBARS levels up to 38.2% only at the end of the study. The *S. maxima* administration in diabetic animals reduced the TBARS levels by15.16%. No obvious reduction in the TBARS levels was observed in *S. maxima* administered normal rats. The Glibenclamide administered normal rats showed a reduction in TBARS level by 16.57% (Figure 1G).

Our study identified the potential role of *S. maxima* in reducing the blood glucose level by addressing various etiologies like oxidative stress and dyslipidemia.

#### 3.1.4 Changes of body weight

Before streptozotocin induction, all the rats had a marginal difference in their body mass. The streptozotocin induction in rats reduced the body mass by 33.37%. The *S. maxima* therapy in rats enhanced the body mass as compared to the diabetic control group.

### 3.2. Lipid content of *S. maxima*

The lipids of *S. maxima* were identified by Gas chromatography mass spectrometry, and the data was interpreted from MS-library (NIST or WILEY 09). Our study revealed, *S. maxima* contained lipids ranging from C_4_-C_22_ chain lengths. The total lipid content of *S. maxima* was found to be 32.5µg/g dry weight. The *S. maxima* lipid substance showed mass fragments m/z 44, 56, 74 which are identified to the characteristic mass peaks of C_5_H_11_NO_2_. These molecules were 1.29% of total lipid content. (Figure 2b).*S. maxima* showed mass spectra peaks at m/z 44, 69, 74, 86 which resembled the mass peaks of C_6_H_13_NO_2._ This C_6_ lipid was identified in *S. maxima* at a level of 1.04% of the total lipids. (Figure 2c). Other mass spectra peaks observed at m/z 53, 63, 81, 110 which are the characteristic peaks of the molecule C_6_H_6_O_2_ (Figure 2d). The C_6_ lipids constituted 17.7% of total lipid content of *S. maxima*. *S. maxima* showed mass spectra peaks at m/z 41, 79, 93 which resembled the mass peaks of C_9_H_16_O. The C_9_ lipid was found in *S. maxima* at a level of 0.68% (Figure 2e).

The C_10_ molecule showed 16.1% of total lipid content (Figure 2f-j). The C_10_ molecules had a molecular weight of 152,138,156,198,148 respectively. *S. maxima* showed mass spectra peaks which resembled the mass peaks of C_11_H_18_ (Figure 2k). *S. maxima* showed mass spectra peaks at m/z 41,69,79,136,149,163 which resembled the mass spectra of C_13_H_22_ (Figure 2l).The C_13_ molecule was 0.093% of total lipid content of *S. maxima*. The C_15_ molecule showed 30.3% of total lipid content and had a molecular weight of 224 (Figure 2m-p). The C15 molecule was found to have molecular peaks m/z 41, 69, 79, 109, 161 which resembled the mass peaks of C_15_H_24_O. The *S. maxima* also showed mass peaks at m/z 40, 41, 55, 69, 79, 120, 133, 161, 175, 189, 204 which resembled the molecule C_15_H_24_. The *S. maxima* also contained C_15_H_26_O with characteristic mass peaks at m/z 41, 53, 79, 91, 185 while C_15_H_32_ showed characteristic mass peaks at m/z 41, 43, 57, 71, 85, 99. The *S. maxima* also showed the presence of C_16_H_32_O_2_ with its characteristic mass peaks at m/z 41, 43, 73, 85, 115, 129, 171, 185, 213, 227, 256. The C_20_ lipids were 7.5% while the C22 lipids were 33.7% of total lipid content of *S. maxima*; The C_20_ lipid showed a molecular peaks at m/z 41, 56 73, 85, 98, 116, 129, 143, 157, 171, 181, 213, 239, 257, 312 and had a close similarity with C_20_H_40_O_2_ (Figure 2q). The C_22_ lipid showed a molecularpe aks 257, 129, 98, 239,213,185, 171, 129, 98, 56 and had a close similarity with C_22_H_44_O_2_. The another molecule had mass peaks at m/z 77, 91, 105, 115, 152, 165, 181, 209 and had a close similarity with C_22_H_22_O (Figure 2r).

### 3.6. Molecular Docking of the Identified Lipids with PPAR-γ and AMPK

The PPAR-γ agonists pose a specific region for several metabolic functions such as insulin sensitivity, glucose uptake through GLUT-4. All the identified lipids of *S. maxima* and their homologies were screened for its interaction with PPAR-γ. We have identified that C_5_, C_6,_ C_11_ and C_18_ class of lipids had the tendency to dock with PPAR-γ. The PPAR-γ molecule and ligand interaction of C5, C6, C11 and C18 molecules have been depicted in the Figures (Fig 3-4). The binding C_5_ molecules CID 4727 had the highest binding affinity (−6.65 k cal) with PPAR-γ. The docking results showed a characteristic binding with Lys 275, Glu 259, Leu 255, Arg 280. The other C_5_ molecules (CID 83693) had a binding energy of −5.7 k cal, and a characteristic binding affinity with His 266, Glu 259, Leu 255, Arg 280. The C_5_ molecule (CID 450622) had a binding energy of −4.95 k cal and had a characteristic binding with Arg 280, Gln 283, His 266, Glu 259. Whereas the molecule CID 11075314 had a binding energy of −5.09 kcal and had a characteristic binding affinity with Arg 280, His 266, Gln 283, Glu 259. The C_5_ molecule (CID 17750942) had a binding energy of −4.03 kcal and had a characteristic binding with Met 252. The molecule CID 2796 possessed a binding energy of −4.96 kcal and had a characteristic binding with Glu 259, Arg 280 whereas CID 16421 had a binding energy of –5.57 kcal and a characteristic binding with lys 275. The C_6_ molecule CID 289 showed binding energy of −5.59k cal and had a characteristic binding with Lys 265, Glu 259, Asp 260. The CID 3610 possessed a binding energy of −4.9 kcal and had a characteristic binding with Arg 280, Phe 264. The molecule CID 4534 possessed a binding energy of −5.07 kcal and had binding affinity with Glu 259, Phe 264. The C6 molecules CID 5816 also had a binding energy of −5.76 Kcal and showed binding affinity with Asp 360,Arg 280,Glu 259. The C_6_ molecule CID 445154 possessed the maximum binding energy of −5.96 Kcal and showed a characteristic binding with Ile 262, Glu 259, Ile 281. The C_11_ lipid molecule possessed a binding energy of −3.17 kcal and a characteristic binding with Glu 259 (Table 6). The study indicated that C_5_,C_6_ and C_11_ lipids of *S. maxima* are capable of binding with the pockets of PPAR-γ (Figure 4 A-C).

**Figure 3.**
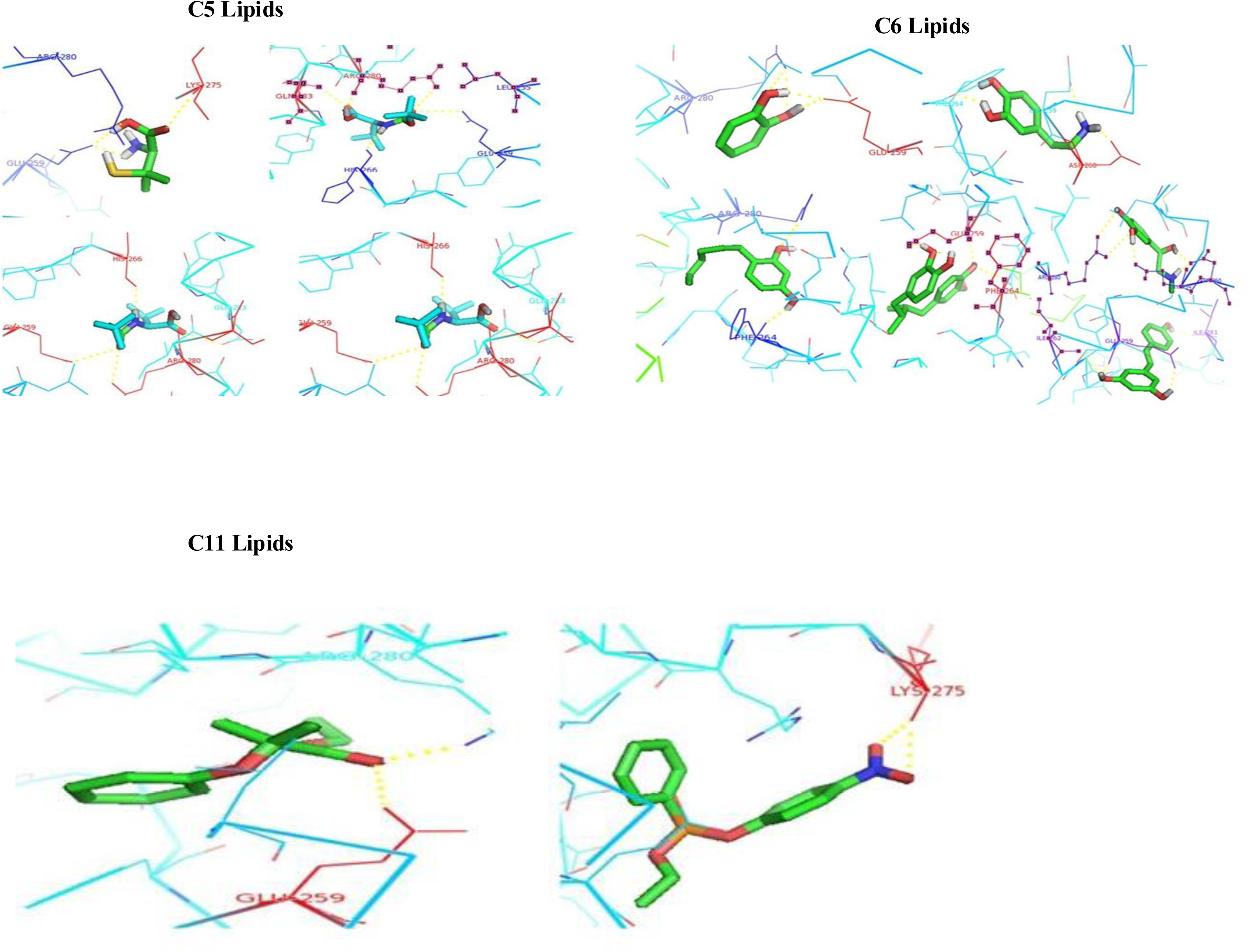
In silico Docking of Bioactive Lipids from *Spirulina maxima* with PPAR-Ɣ

**Figure 4.**
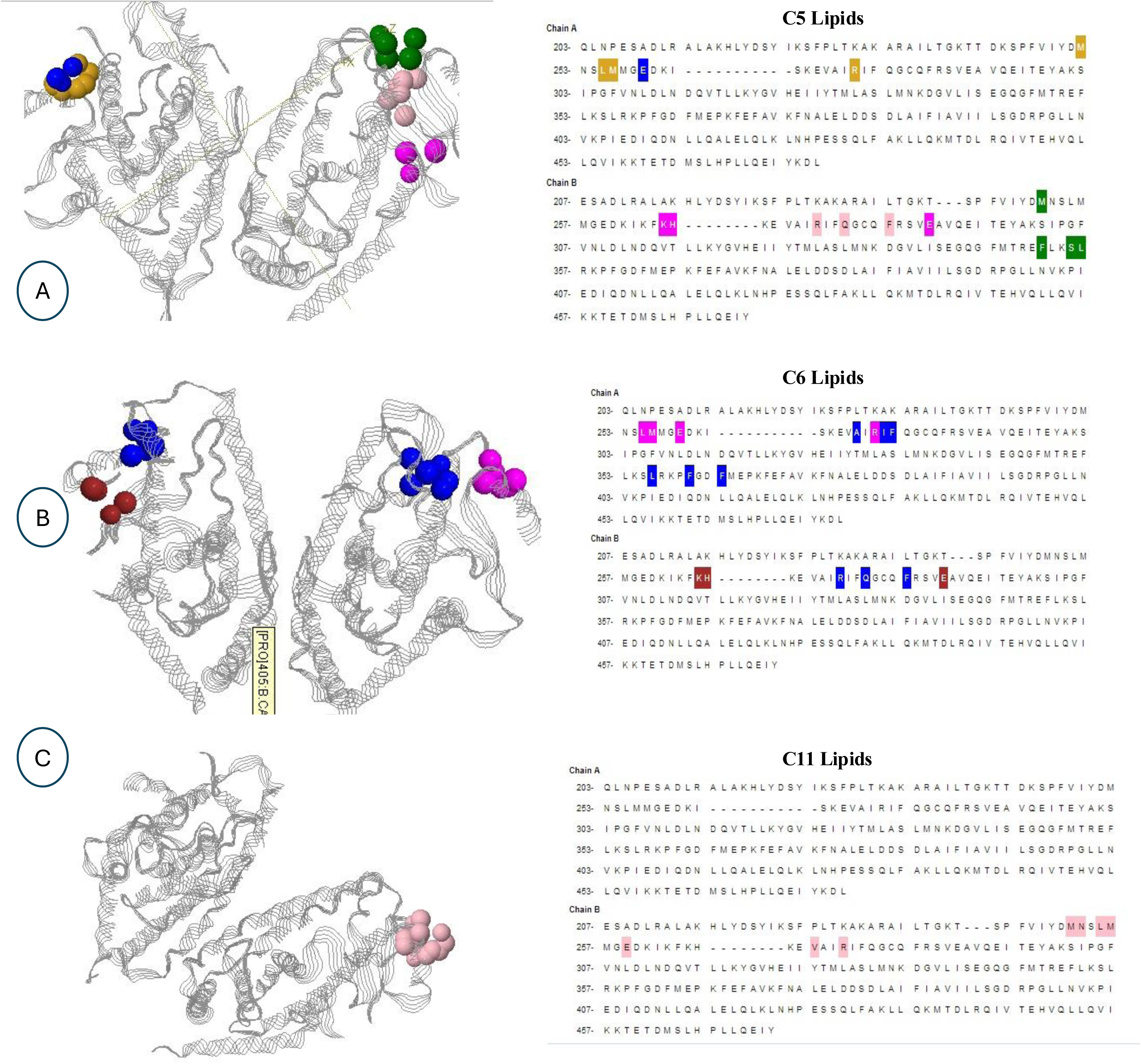
Molecular docking analysis of Spirulina maxima lipids showing binding sites and interacting residues within PPAR-γ

Our study also revealed that C5, C6, C10, C11 and C18 lipids have identical binding sites on AMPK that is Arg 69, Ser 225, Arg 151 (Table 5). It has been evident from thee molecular docking analysis that *S. maxima* lipids doesn’t exactly bind on activation site of AMPK, however it bind exactly on the other functional domains which facilitates the autoinhibition of AMPK activity.

**Table 5:** Binding Affinity and Amino Acid Interactions of AMPK with Spirulina Lipids Identified by Molecular Docking.

|  |  |  |
| --- | --- | --- |
| Docking of AMPK with C5 molecules |  |  |
| CID | Binding energy | Amino acids |
| 11075314 | -6.65 | Arg 69, Ser 225, Arg 157 |
| 9882671 | -5.7 | Asp 316, ser 315, ser 225, Al 226 |
| 6971276 | -4.95 | Ser 225, Arg 298, Ser 241 |
| 450622 | -5.09 | Arg 151, Arg 69 |
| 94744 | -4.90 | Arg 69, Ser 241, ser 225 |
| Docking of AMPK with C6 molecules |  |  |
| CID | Binding energy | Amino acids |
| 681 | -5.55 | Lys 148, His 168, Asp 316 |
| 785 | -5.55 | Lys 169, Val 292 |
| 5054 | -5.24 | Val 292, Val 296 |
| 14537 | -5.69 | Leu 314, Ser 313, Ser 315 |
| 16043 | -5.14 | His 168, Lys 169,His 197 |
| Docking of AMPK with C10 molecules |  |  |
| CID | Binding energy | Amino acids |
| 2083 | -5.73 | Ser 313, Gln 314, leu 145 |
| 4260 | -5.27 | Val 224, Ser 225, Asp 316, Arg 298 |
| 6450 | -5.27 | Ile 145 |
| 6620 | -6.14 | Arg 298, lys 242 |
| 9339 | -9.49 | Ser 313, ser 225 |
| 5i678 | -5.74 | Ser 225, val 224, Thr 199 |
| Docking of AMPK with C11 molecules |  |  |
| CID | Binding energy | Amino acids |
| 559050 | -5.35 | Arg 151 |
| 1940914 | -4.23 | His 150 |
| 3457348 | -5.75 | Ser 225, Ala 226 |
| 10889009 | -4.84 | Lys 169, Arg 293, Ser 225 |
| Docking of AMPK with C18 molecules |  |  |
| CID | Binding energy | Amino acids |
| 11005 | -5.74 | Leu 166, Arg 69, Arg 151 |
| 16213483 | -7.1 | Arg 69, |
| 16213484 | -6.58 | Arg 69, Arg 151 |
| 53378100 | -6.29 | Arg 69, Arg 151 |

## 4.0 Discussion

Diabetes is a major metabolic disorder affecting nearly 10% of the population all over the world. It results from a shortage or lack of insulin secretion or reduced sensitivity of the tissue to insulin. Diabetes is a metabolic disorder of carbohydrate and lipid metabolism(s) associated with oxidative stress. There are sporadic reports on the anti-diabetic activity of *Spirulina sp.* (Mani et al., 1998; Layam et al., 2006). Our earlier investigation has also reported the potential antidiabetic effect of *S. maxima* in fructose induced hyperglycemia and hypertriglyceridemia in rats (Jarouliya et al., 2012). In the present study, we attempted to figure out whether *S. maxima* administration had its potential antihyperglycemic effect in Streptozotacin induced diabetes mellitus. The study was also attempted to characterize the metabolites of *S. maxima* through GCMS and with the help of molecular docking approaches attempts have been made to screen the molecules, which function as PPAR-γ agonists, at present considered as the safest drug target for antidiabetic therapy (Pedro Iglesias and Juan J Díez 2006)

Attempts were made to optimize the *S. maxima* dosage for glycemic regulation in normal rats through oral glucose tolerance test. Glucose tolerance test performed after oral administration of *S. maxima* showed a marked glucose lowering effect at an increasing order of dosage ranging from 100mg/kg to 500 mg/kg. Hence 500mg/kg dosage of *S. maxima* was administered in STZ induced diabetic animals. Streptozotocin has a property of selective destruction of pancreatic β-cells through oxidative stress generation and thereby create hyperinsulinemic condition. Moreover this condition is also accompanied by body changes like reduction in body weight. The reduction in the body weight could be due to the heavy protein loss, as a result of STZ mediated β-cell death and unavailability of glucose to the cells.

Oral administration of *S. maxima* over a period of 15 days elevated the body weight in comparison to the diabetic control. The study reveals the fact that *S. maxima* being a nutrient rich diet replenish the protein loss and enhance the body weight. This evidence was in agreement with the studies of Layam et al., (2006), which stated that Spirulina administration minimized the body weight loss which is a characteristic feature of STZ induced diabetes mellitus. *S. maxima* administered in diabetic animals for 15 days significantly lowered the FBG levels. The blood glucose reduction was at par with that of potential antidiabetic drug Glibenclamide. Whereas the Spirulina (15mg/kg) administration for 30 days showed a significant blood glucose lowering effect in STZ induced diabetic animals (Layam and Reddy 2006). Here we assume that increasing the *S. maxima* dosage up to 500mg/kg might have enhanced the speedy recovery of β-cells. Our *in-silico* showed *S. maxima* lipids have a characteristic binding with AMPK molecule. The blood glucose regulation could attributed to ampk activation.

### *S. maxima* beneficial on Dyslipidemia

The lipid levels were also modulated after STZ induction, lipid abnormalities in the pancreatic tissue was already established (Levy et al., 1988). In the present study, elevated levels of cholesterol, triglycerides, VLDL, and LDL were observed. *S. maxima* administration lowered the cholesterol, triglycerides, VLDL, and LDL levels. The HDL cholesterol levels also showed notable elevation after *S. maxima* administration. The studies of Mani et al., (2002); de Caire et al., (1995) reported the increased HDL cholesterol levels after *Spirulina* consumption and furthermore, identified γ-Linolenic acid to be responsible for the lipid regulatory mechanisms. These γ-Linolenic acid and other PUFA again serve as natural PPAR-γ agonists (Lai et al., 2013). Our *in-silico* study also indicated C_5_, C_6_ and C_11_ lipids as a potent ligand for PPAR-γ. The lipid lowering effect of *S. maxima* could be attributed to the presence of certain lipids which may participate in the activation of PPAR-γ molecule and regulate the lipid metabolism.

### *S. maxima* beneficial on an Oxidative stress

One of the mechanisms of STZ induced pancreatic β-cell damage is mediated through the generation of free radicals. In the present study, it was observed that STZ induction significantly the levels of GSH, SOD and Catalase while elevated the TBARS levels. The data indicated elevation in the levels of GSH, SOD and Catalase after *S. maxima* therapy while the TBARS levels significantly decline. The increase in the levels of antioxidant enzymes could be attributed to the presence of carotenoids phenolic content and phycocyanin present in *S. maxima*. Estrada et al., (2001) reported the antioxidant property of the protean extract of *S*. *platensis.* Miranda et al., (1998) also reported the antioxidant protection of *Spirulina* extracts *in-vitro* and *in-vivo* systems. The present study concluded the antioxidant potential role of *Spirulina maxima* in STZ induced diabetes in rats.

#### Characterization of *S. maxima* lipids

The cyanobacterial lipids are often modulated by environmental changes such as temperature, nutrient stress (Sato and Murata 1982). In the present study, we observed C_16_, C_20_ and C_22_ lipids in *S. maxima* cultivated in Airlift photobioreactor under controlled temperature 30^0^c. The C_22_ lipids contributed the major part of the total lipid content of *S. maxima*. The C_16_ and C_20_ fatty acids were constituted less than (<10%) of total fatty acid, whereas the C_18_ fatty acids remained undetectable. I was hypothesized from the study that temperature and light may play an important role in enhancing the C_18_ lipids of *S. maxima*. Several other studies also reported, low temperature might facilitate the production of C_18_ lipids in *Spirulina maxima*. This study is in agreement with the fact that chain length of fatty acids and the degree of unsaturation varied with the temperature (Patterson 1970; Klenschmidt 1970;Brown and Rose 1969). Apart from the fatty acids, *S. maxima* also contained lipids with varying chain length and concentration. The C_15_, C_10_ and C_6_ lipids constituted above (>10%) of total lipid content of *S. maxima*. The C_5_ and C_11_ lipids constituted less than (<10%) of total lipid content of *S. maxima*. The C_8_, C_9_ and C_13_ lipids were found to be less than 1% of total lipids present in *S. maxima.* It was found that the lipid content of *S. maxima* is not affected by the temperature and culture conditions. This study was in agreement withthe studies of Quoc and Dubacq (1997) revealed no changes in chain length of fatty acids of *S. platensis* with growth temperature or in linoleate-supplemented media. The study concluded the presence of several lipid molecules with varying chain length ranging from C_5_ to C_22_ lipids; however, the bioactivity of these lipids remain obscure. Hence the insilico molecular docking approach was used to screen the lipid molecules of *S. maxima* for its functional bioactivity against potent antidiabetic targets PPAR-γ and AMPK.

### Docking of Spirulina lipids with PPAR-γ

PPARs belonging to the nuclear receptor have been identified as the safest antidiabetic target. Among these PPARs, PPAR-γ has dragged especially attention, as it involves in the multiple mechanism of carbohydrate metabolism, lipid metabolism. Hence it plays an important role in insulin resistance, dyslipidemia and inflammation. PPAR-γ function via activation through ligands and regulates the expression of genes involved in glucose and lipid metabolism. PPAR-γ activation can influence insulin signaling at various steps in these pathways, resulting in improved whole body insulin sensitivity and enhanced glucose and lipid metabolism. In the present study, we have identified that some of the lipid molecules (C_5_, C_6_, C_11_, and C_18_) are capable of binding with the PPAR-γ molecule. We identified that all the lipid molecules had specific binding in the ligand binding regions of PPAR-γ (Regions 241-421), the most common binding sites for all these ligands were Glu 259 and Arg 280. The C_5_ and the C_6_ molecules had higher binding sites in the ligand binding regions. In our study, streptozotocin was used to develop type 1 diabetes mellitus. Hence we believe that the speedy recovery of the β-cells damage after *S. maxima* therapy could be due to the activation of PPAR-γ, which facilitate the glucose uptake and reduce the glucotoxicity in pancreas.

AMPK is the power regulator of the cells. The AMPK molecule senses the blood glucose levels and serve as a regulator of several catabolic and anabolic pathways of carbohydrate, lipid and protein metabolism. Our study showed C_5_, C_6_,C_11_ have a characteristic binding affinity in the gamma subunit of AMPK. The blood glucose regulation could be due to the binding of these lipids with the AMPK.

## Conclusions

From the above the study it was concluded that *S. maxima* administration regulated the blood glucose homeostasis in diabetic rats and also regulated the lipid metabolisms. The *S. maxima* administration also reduced the oxidative stress developed in diabetic condition by enhancing the activities of the counteracting antioxidant enzyme. The *in-silico* docking concluded that some of the lipid molecules of *S. maxima* has an effective binding affinities with AMPK and PPAR-γ.

